# Electrospinning of pure, native, cross-linker free self-supported collagen membrane

**DOI:** 10.1101/616946

**Authors:** Dounia Dems, Julien Rodrigues da Silva, Christophe Hélary, Frank Wien, Marie-Claire Schanne-Klein, Christel Laberty-Robert, Natacha Krins, Carole Aimé

## Abstract

Electrospinning (ES) is an extremely promising method for the preparation of self-supported membranes for tissue engineering by mimicking the 3D fibrillar structure of the extracellular matrix. Conflicting results about collagen ES in the literature concern the conditions of collagen solubilization to improve electrospinnability, and the use of co-polymers and chemical cross-linkers to stabilize the structure of collagen membranes. Here we report for the first time (1) the ES of pure and native collagen into a self-supported membrane in absence of polymer support and (2) the preservation of the membrane integrity in hydrated media in absence of crosslinker. We use a multiscale approach to characterize collagen native structure at the molecular level using synchrotron radiation circular dichroism, and to investigate collagen hierarchical organization within the self-supported membrane using electron and multiphoton microscopies. Finally, we show that the membranes are perfectly suited for cell adhesion and spreading, making very promising candidates for the development of advanced biomaterials.

## 1. Introduction

Mimicking the structure of the extracellular matrix (ECM) is of primary importance in tissue engineering to provide the adequate tridimensional (3D) environment and biological cues for cell organization and function. Electrospinning (ES) has become increasingly popular as a cost effective technology to produce nanofiber scaffolds for tissue engineering.^1^ In the ES process, high voltage is applied to a polymer solution that is ejected through a needle towards a grounded collector as dried fibers. Upon voltage application, the liquid droplet deforms into a cone, the “Taylor cone”, and emits a charged jet of liquid. The charged jet is then elongated in a whipping process and moves towards a grounded collector. ES allows to produce fibers with tunable diameter (smaller than 1 μm) and orientation. Biopolymers are the most relevant candidates to be electrospun to mimic the ECM as they bear the biochemical cues necessary for cell attachment and proliferation. As major component of ECM, collagen is of great interest for biomedical applications. The structure of the collagen network consisting of fibrils (diameter from 20 to 500 nm) is crucial for cell attachment, proliferation, and differentiation. ES of collagen remains challenging. Collagen solutions do not have the viscoelastic response necessary for jet stabilization partly due to the stability of collagen tertiary structure.^2^ As a result, denaturing fluoroalcohols are often used, such as 1,1,1,3,3,3-hexafluoro-2-propanol^3–8^ and 2,2,2-trifluoroethanol,^6^ allowing stretching and rearrangement of collagen. Since then, the preservation of the native state of collagen has continuously been a subject of debate.^9^ Several works evidence that collagen completely unfolds in fluorinated solvents. Efforts have been made to find alternative solvents, such as phosphate buffer saline (PBS)/ethanol^10^ or acetic acid/DMSO^11^ mixtures. All of them involve the use of chemical crosslinkers (glutaraldehyde^3,12^ or carbodiimide^10,11,13,14^ to prevent the membrane from falling apart in hydrated environment. However, the use of crosslinkers alters the fibrillogenesis of collagen and the biocompatibility of the materials.^15^ Original approaches have been developed using a polymer support, where collagen is electrospun together with a synthetic polymer (polycaprolactone,^5,6,16,17^ polyethylene oxide,^14,18^ polyvinyl-pyrrolidone^19^) eventually washed out after crosslinking, or with a biopolymer^20^ including elastin,^14^ chitosan,^7^ or gelatin.^21^

In this work, we report for the first time the ES of pure collagen from rat tail tendons into a self-supported membrane in absence of polymer support (Fig. 1A). We use an acidic/alcoholic mixture of non-denaturing solvents that fully preserves the native triple helical structure of collagen. In addition, we show how to efficiently stabilize the membrane structure for further applications in biological hydrated environment in absence of crosslinkers by inducing collagen pre-fibrillation in basic conditions. We use a multiscale approach, from the molecular scale, to characterize the triple helix of collagen using synchrotron radiation-circular dichroism (SRCD), to the nanometer and micrometer scales to probe collagen organization within the membrane using field emission scanning electron microscopy in a cryo mode (Cryo-FE-SEM) and multiphoton microscopy based on second harmonic generation (SHG). This approach demonstrates that we can provide an easy procedure to handle biomaterials made of pure electrospun collagen fibers exhibiting the hierarchical organization of collagen observed in living tissues, which allows cell adhesion and spreading.

**Figure 1.**
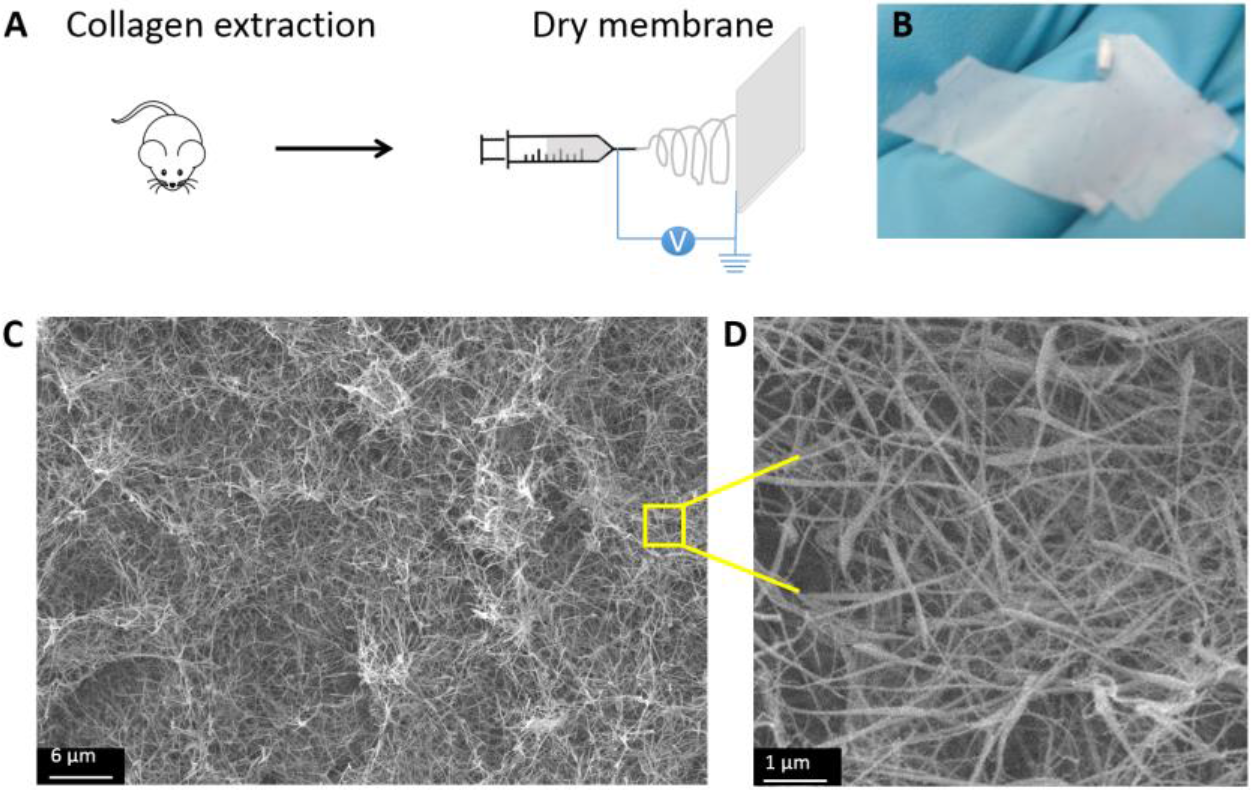
(A) Scheme of collagen ES. (B) Photo, (C,D) SEM images of a self-supported membrane ([collagen] = 1.7 mg.mL-1 in 30 mM HCl:EtOH (25:75 v/v)).

## 2. Results and Discussion

### 2.1. A self-supported membrane of pure collagen

Many parameters can be adjusted during the ES process to get a stable jet and ensure nanofiber formation. This concerns the ES process by itself (potential, flow rate, needle to collector distance, collector shape and composition), the solution (viscosity, conductivity, surface tension, concentration), as well as external parameters (temperature, humidity). Among those, the appropriate selection of a solvent system is a prerequisite for successful collagen ES, while avoiding denaturation. We focused on the solvent selection and ultimately adapted the ES parameters, working in a home-made chamber flushed with dry air and with a copper collector.

Collagen triple helix is stable in solution in acidic conditions. Hydrochloric acid (3 mM HCl, pH 2.5) was preferred over acetic acid routinely used for collagen due to its higher evaporation rate. Further improvement of electrospinnability was reached by adding ethanol (1:1 vol) to the collagen solution (7 mg.mL^−1^). In these conditions, only large beads could be observed under conventional SEM (Fig. S1A,B). The formation of beads can be attributed to jet instability, which can be reduced upon increasing solution viscosity. Solution conductivity was also found to influence fiber formation.^16,22^ To comply with those requirements, the ethanol content was increased up to 75% vol, increasing the viscosity of the solution, without preventing ES. In those conditions, to prevent the formation of a gel, collagen concentration was decreased from 7 to 4 mg.mL^−1^. This optimization step already showed efficient elongation of the beads (Fig. S1C-D). Decreasing further collagen concentration to 2 mg.mL^−1^ allowed to obtain a self-supported membrane with a thickness of 40 µm, but exhibiting short fibers still ending with large beads (Fig. S1E-F). It is worth noting that the working regime, 0.12 wt% collagen, is particularly low when compared to the polymer concentrations commonly used for electrospinning (6-10 wt%). This is attributed to the fact that collagen is a highly charged polymer. Further improvements were obtained by increasing HCl concentration to 30 mM. In those conditions, randomly oriented and interconnected fibers with diameter in the nanometer range (58 ± 10 nm) were produced, where the number of beads could largely be decreased and exhibiting an elongated shape (Fig. 1 B-D). Increasing HCl concentration up to 300 mM did not improve further the fiber morphology. By varying the nature of the acid used, the ethanol content and the concentration of collagen, we successfully set the conditions to comply with solvent evaporation, solution viscosity and conductivity necessary for collagen electrospinning in absence of denaturing solvent. The macroscopic structure of those networks is described in more details in Fig. S2. In the following, collagen has been electrospun at a concentration of 1.7 mg.ml^−1^ in 30 mM HCl and 75% vol. ethanol.

### 2.2. Preserving the native triple helix structure of collagen

Whether collagen may be denaturated remains an open question in literature. This can involve both the solvent used for improving the electrospinnability of the collagen or the spinning process by itself and the high voltage applied. SRCD has been used to investigate collagen secondary structure after ES. To this aim, self-supported membranes were dissolved in 3 mM HCl. A large negative band at 198 nm and a small positive band at 223 nm were observed at 20°C (Fig. 2A solid line). This corresponds to the spectral signature of the collagen triple helix after extraction and before ES, and characteristic of the polyproline II (PPII) conformation of collagen triple helix (Fig. 2A, dotted line). This CD spectrum strikingly deviates from previous studies that showed collagen denaturation,^8,23^ and evidences that collagen fully preserves its native state after ES. Then, the melting temperature (T_m_) of the soluble triple helix was measured by thermal denaturation with increasing temperature up to 74°C in steps of 3°C. As expected, the intensity of both bands decreases with the temperature increase (Fig. 2B from dark blue to dark red). This is attributed to the increased fraction of unfolded structures upon thermal denaturation. T_m_ was determined by plotting the ellipticity at maxima (198 and 223 nm) as a function of temperature using the sigmoidal fit in “Igor Pro” spectral analysis program.

**Figure 2.**
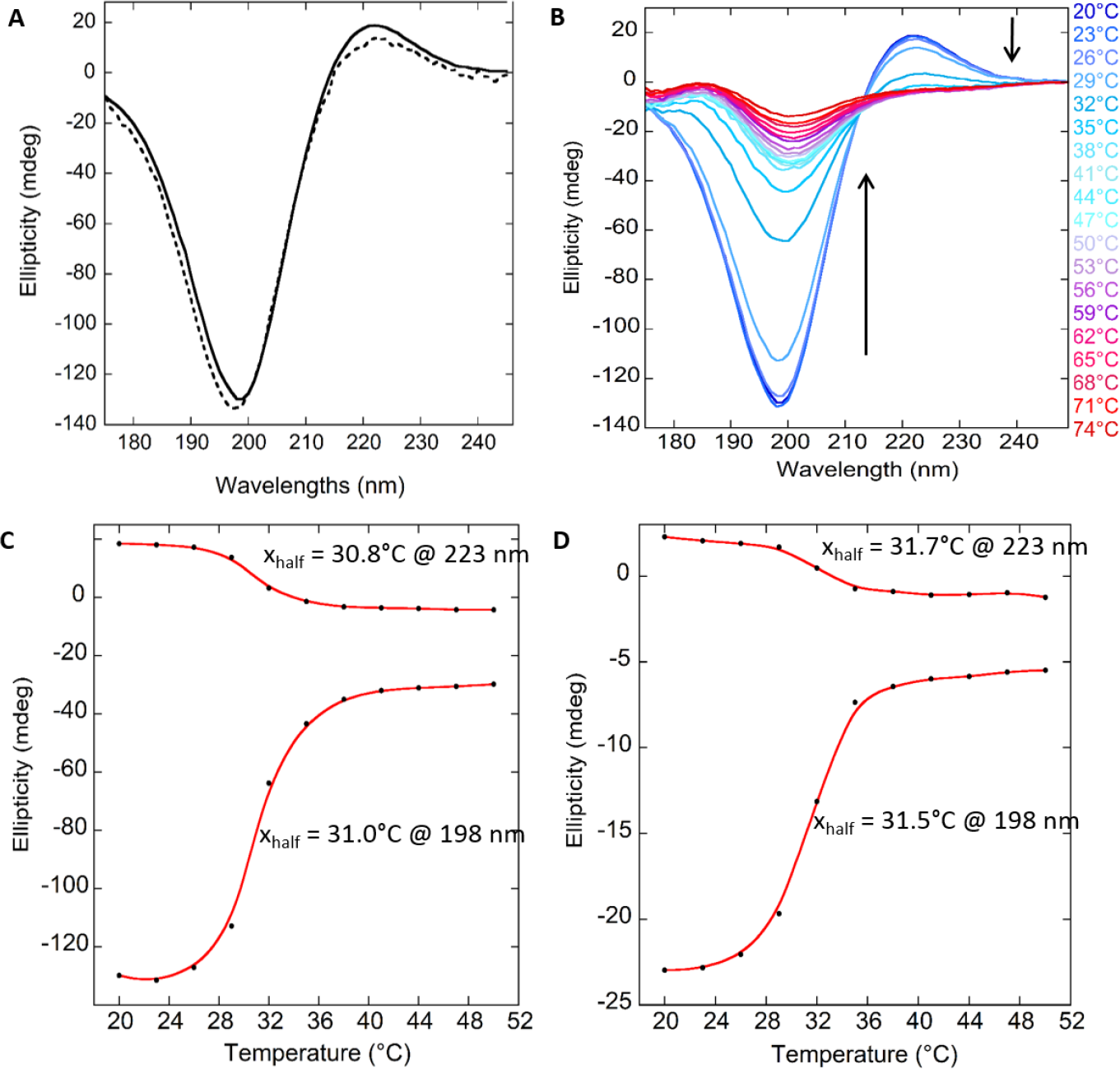
SRCD spectra of (A) collagen in 3 mM HCl before (dotted line) and after (solid line) ES, and of (B) thermal unfolding of ES collagen from 10 to 74 °C in steps of 3°C. (C-D) Ellipticity as a function of temperature: Tm at 198 and 223 nm for ES and native collagen respectively.

After ES, the T_m_ was found to be around 31°C at 198 nm and 30.8°C at 223 nm (Fig. 2C), in agreement with previously reported values for native collagen.^9^ Above 31°C, the structure is dominated by unordered conformations, and collagen completely unfolds above 40°C. To estimate the impact of the ES process, we compared the thermal denaturation of the electrospun triple helix with that of native collagen after extraction. Figure 2D shows that T_m_ of pure collagen is similar to the one of the ES membrane (31.5°C at 198 nm and 31.7°C at 223 nm). This shows that the fraction of PPII at room temperature in both samples is the same and further confirms the preservation of the native state of collagen after ES.

### 2.3. Stabilization of the scaffold structure in hydrated conditions

In a context of biomedical applications, a key question to address is the stabilization of the membrane structure in a hydrated form. Indeed, when immersing the as-obtained membrane in a hydrated environment (typically cell culture medium), the collagen membrane falls apart. To tackle this issue, works reported in the literature often use chemical cross-linkers that can be detrimental for cell survival, while hampering collagen hierarchical self-assembly. Alternative strategies have to be developed to strengthen the hierarchical structure of collagen. To this aim, we exposed ES membranes to ammonia vapors, to take advantage of the increase in pH to stabilize collagen fibrils. This strategy is known to trigger fibrillogenesis and gelling in collagen solution^24,26^ but has never been adapted to dry materials. Very efficiently, after a 20 minutes exposure of the dry membrane, the macroscopic structure of membranes was unchanged but they could be stored in culture medium over months. SHG and cryo-FE-SEM were used to investigate the hierarchical structure of the ES membrane in the course of the stabilization and hydration. We compared (A) the dry membrane, as obtained after ES, (B) the membrane after 20 min exposure to NH_3_ vapors and (C) the hydrated membrane in cell culture medium (Fig. 3A-C).

#### Second harmonic generation (SHG)

SHG is a coherent process specific for non-centrosymmetric materials. It builds up efficiently in highly anisotropic fibrillar collagen because of the tight unidirectional alignment of peptide bonds along the collagen triple helix and within fibrils both in isolated collagen fibrils^27^ and fibrillary networks.^28^ This makes SHG microscopy the gold standard technique for 3D characterization of collagen-rich tissues in ambient conditions and without exogeneous labelling.

**Figure 3.**
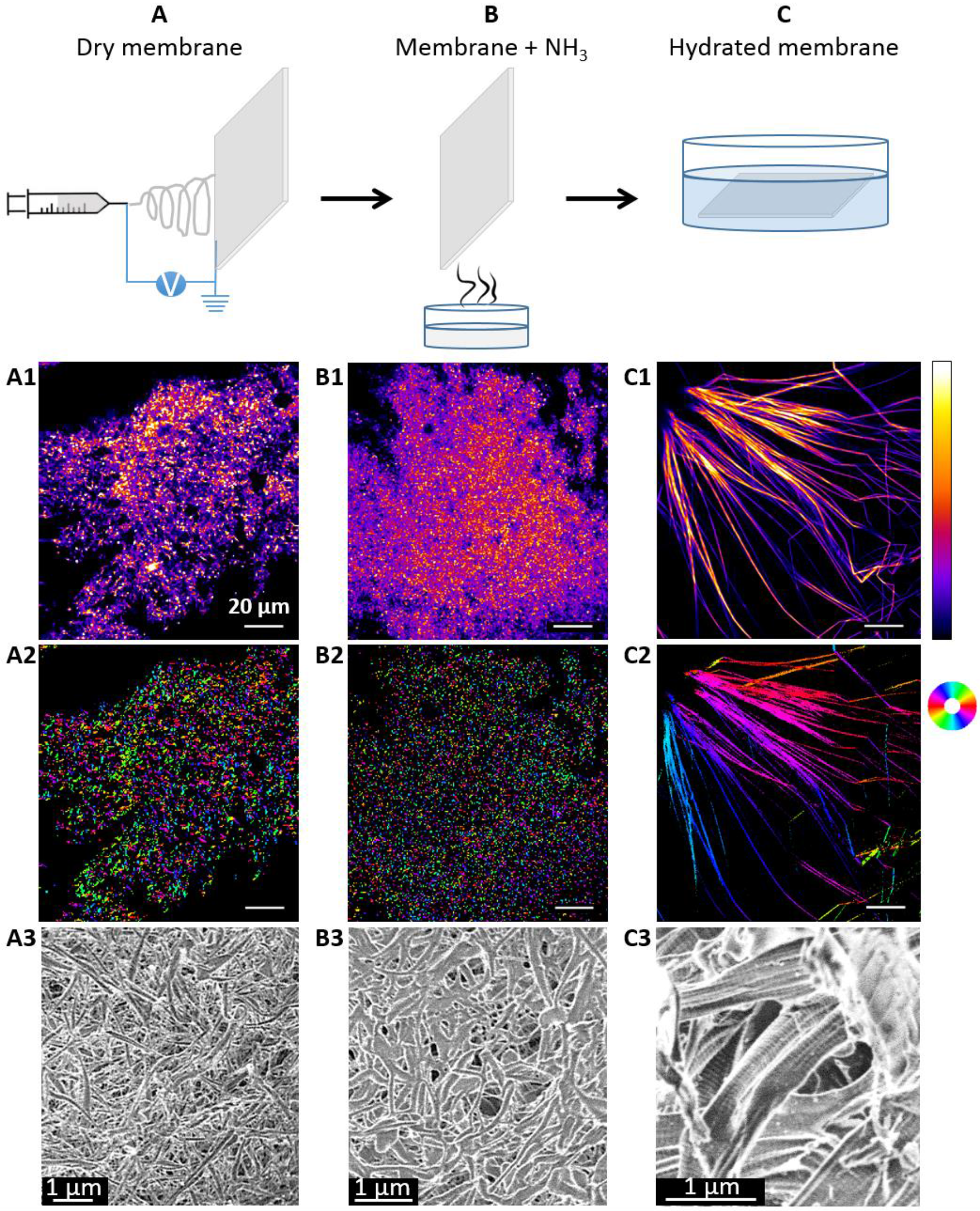
(1) SHG, (2) pSHG and (3) cryo-FE-SEM images of (A) a dry collagen membrane, (B) after exposure to ammonia vapors and (C) rehydration in culture medium. The look-up table (on the right in the A1-C1 line) corresponds to arbitrary units for each SHG image. The orientation in the pSHG image is displayed on the colored circular scale (on the right in the A2-C2 line).

Measurements on dry membranes showed a high SHG signal (Fig. 3A1). This is an additional proof of the native state of collagen given the high specificity of SHG for collagen triple helices over gelatin. In addition, this is indicative of the non-centrosymmetric packing of the triple helices. At this stage, it remains difficult to know whether collagen is fibrillated or not, as SHG is also observed in liquid crystal phases.^29^ In addition, the grainy-like pattern with a myriad of dots indicates the presence of small structures that may be highly entangled, in agreement with SEM observations (Fig. 1C,D). Indeed, if the diameter of the fibrils is small relatively to the focal volume (58 ± 10 nm in diameter for a resolution of typically 0.4 μm (lateral) × 1.7 μm (axial)), many fibrils may be simultaneously observed in one pixel. At the intersection of many fibrils, molecules in the focal volume are not all oriented in the same direction, which results in a centrosymmetric distribution and leads to a local strong decrease of SHG signal (Fig. 4A-B). If the network is highly entangled, SHG extinction is widespread throughout the focal volume leading to a grainy-like image. The structural specificity of SHG images has been further improved by acquiring polarization-resolved images (pSHG), which measures the main orientation of collagen in the imaging plane, in every pixel of the SHG image.^30^ Given that each color codes for a given direction, the presence of dots with many different colors throughout the whole image shows that the highly packed structures of native collagen are randomly distributed with respect to one another (Fig. 3A2). Again, at fibrils intersection, all molecules are not oriented in the same direction, which impedes the determination of a main direction, resulting in black pixels (Fig. 4C-D). After 20 min exposure to NHM_3_ vapors, no variation of the SHG and pSHG profiles could be detected (Fig. 3B1-2). This shows the preservation of the triple helix structure and packing within small entangled structures. No important variation at the scale probed by SHG can be detected despite the very different behaviors of the dry and stabilized membranes when immersed in hydrated environment.

In contrary, SHG observations of the hydrated membrane showed very important variations of the membrane structure that appears made up of large micrometer-long fibrils (Fig. 3C1). This is reminiscent of collagen matrices in dense fibrillated media. pSHG very nicely reveals the unidirectional ordering of the triple helices in each collagen fibril that appears as monocolored structures (Fig. 3C2). In this case, the absence of grainy-like profile is attributed to the fact that collagen reorganization leads to the formation of large structures. Indeed, one of the major features of type I collagen is its ability to self-assemble into highly organized supramolecular structures, which in turn form larger highly oriented fibers and fascicles. This prevents the observation of multiple collagen fibrils with random organization with respect to each other within the focal volume, hence precluding local SHG extinction.

**Figure 4.**
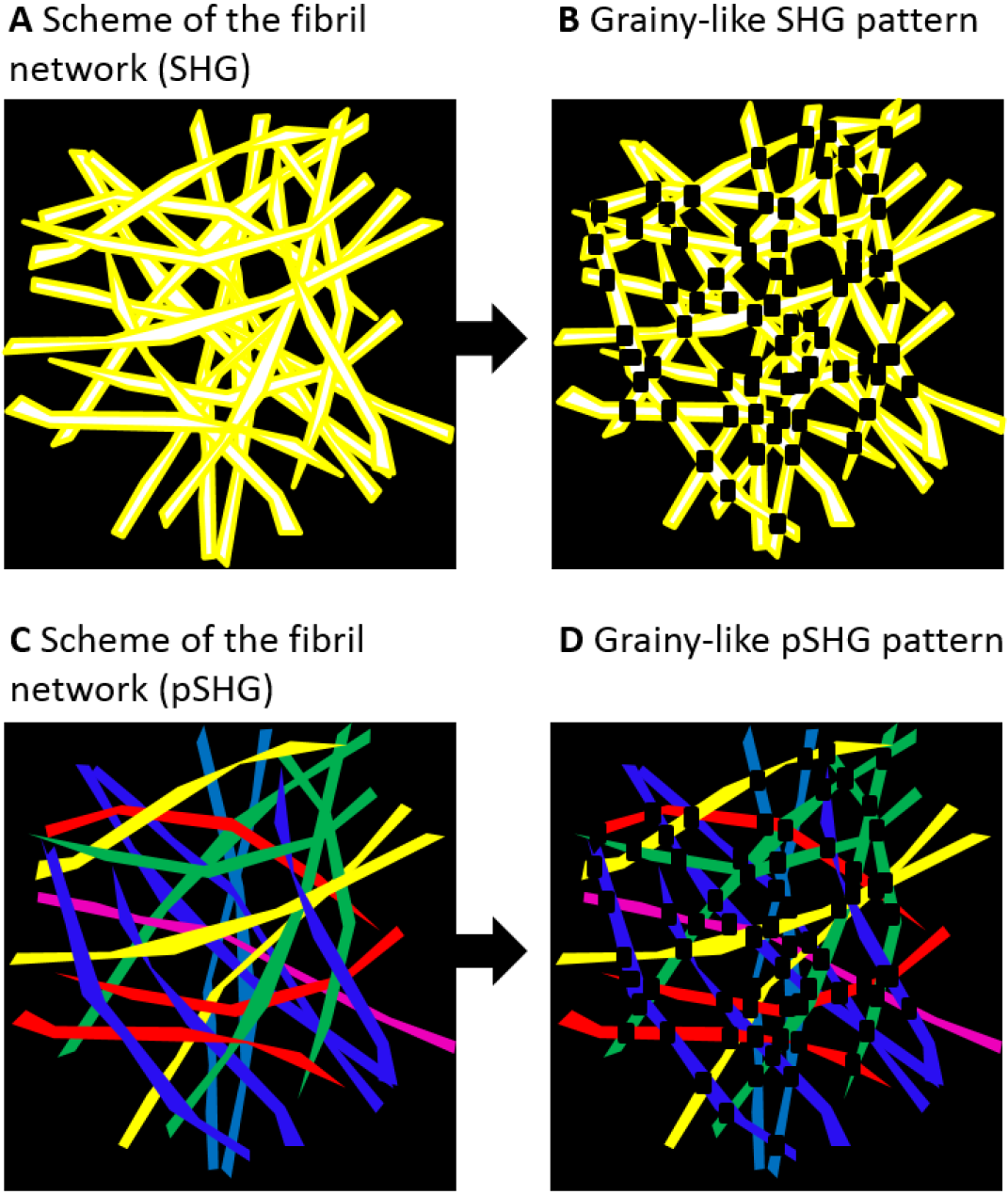
Schematic representation of the local decrease in SHG intensity in highly entangled network of small diameter-fibers. Scheme of the fibril network and observation of the corresponding grainy-like pattern by (A-B) SHG and (C-D) pSHG, showing local strong decreases in SHG and pSHG intensities at the intersection of collagen fibers.

#### Cryo-FE-SEM

Under cryo-FE-SEM that preserves the hydration state of collagen membranes, the dry membrane exhibits the same structure as observed under conventional SEM due to the absence of solvent within the initial biomaterial: randomly entangled fibrils are clearly visible (Fig. 3A3). Once stabilized under NH_3_ vapors, the overall structure of the sample remains unchanged, exhibiting an entangled network but with slightly melted fibrils (Fig. 3B3). This shows that ammonia exposure does not affect the 3D organization of the collagen nanofibers. This is a very interesting alternative to crosslinking strategies usually used to stabilize collagen-based membranes. Unlike previous works, in this case no gelation process is induced: ammonia exposure allows for the membrane storage in pre-fibrillated state. This can be attributed to a very limited extent of fibrillation at the solid-vapor interface hampered by the low amount of water as diffusing medium. After hydration, large fibrils could be observed exhibiting the 67-nm periodic pattern characteristic of collagen organization in living tissues (Fig. 3C3). These fibrils further organized into bundles of tightly aligned fibers exhibiting large width and polydispersity (from 200 nm to 2 µm, with a mean width of 560 ± 290 nm over a population of 61 fascicles). This very nicely confirms the variations of the collagen network observed by SHG/pSHG and again proves the native state of collagen within ES membranes given that non-native collagen is not able to form cross-striated fibrils. Very importantly, this ascertains the possibility to recover the self-assembling processes and hierarchical organization of collagen as observed in living tissues. This is the first time that such organization could be obtained after ES.

### 2.4. Fibroblast adhesion and differentiation

To assess the cytocompatibility and bioactivity of electrospun collagen nanofibers, cell attachment and spreading of normal human dermal fibroblast cells (NHDF) seeded onto the collagen matrix were studied. Immunostaining of actin and nuclei revealed the colonization of the scaffold, where the fibroblasts were spread all over the membrane and exhibited a spindle-like shape morphology with stress fibers clearly visible inside the cellular body (Fig. 5, white arrows). This clearly evidences that ES membranes are perfectly suitable for cell adhesion and spreading.

**Figure 5.**
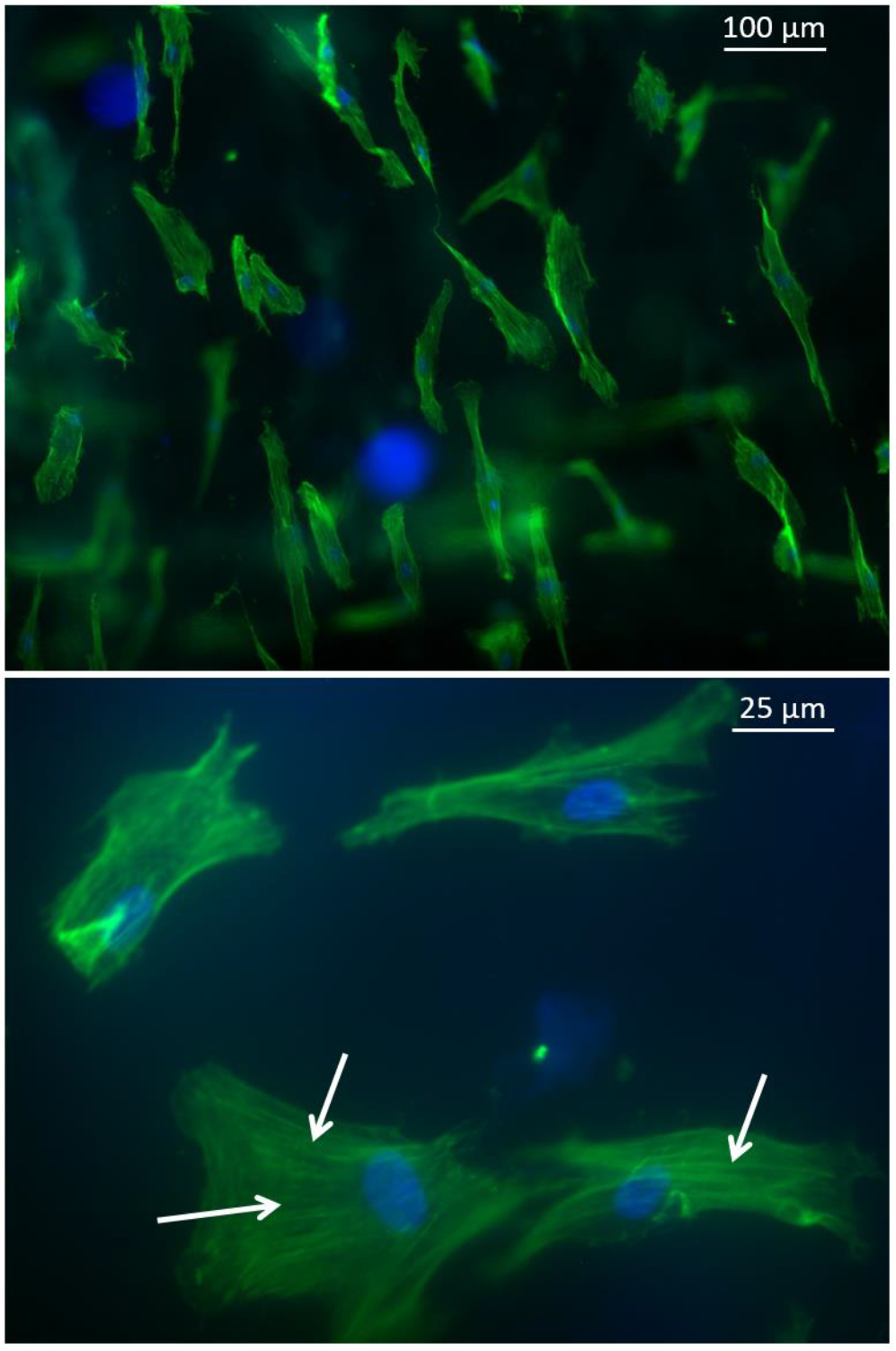
NHDF cultured for 24 h on ES membranes. Actin (green: phalloidin) and nucleus (blue: DAPI) staining. White arrows show stress fibers.

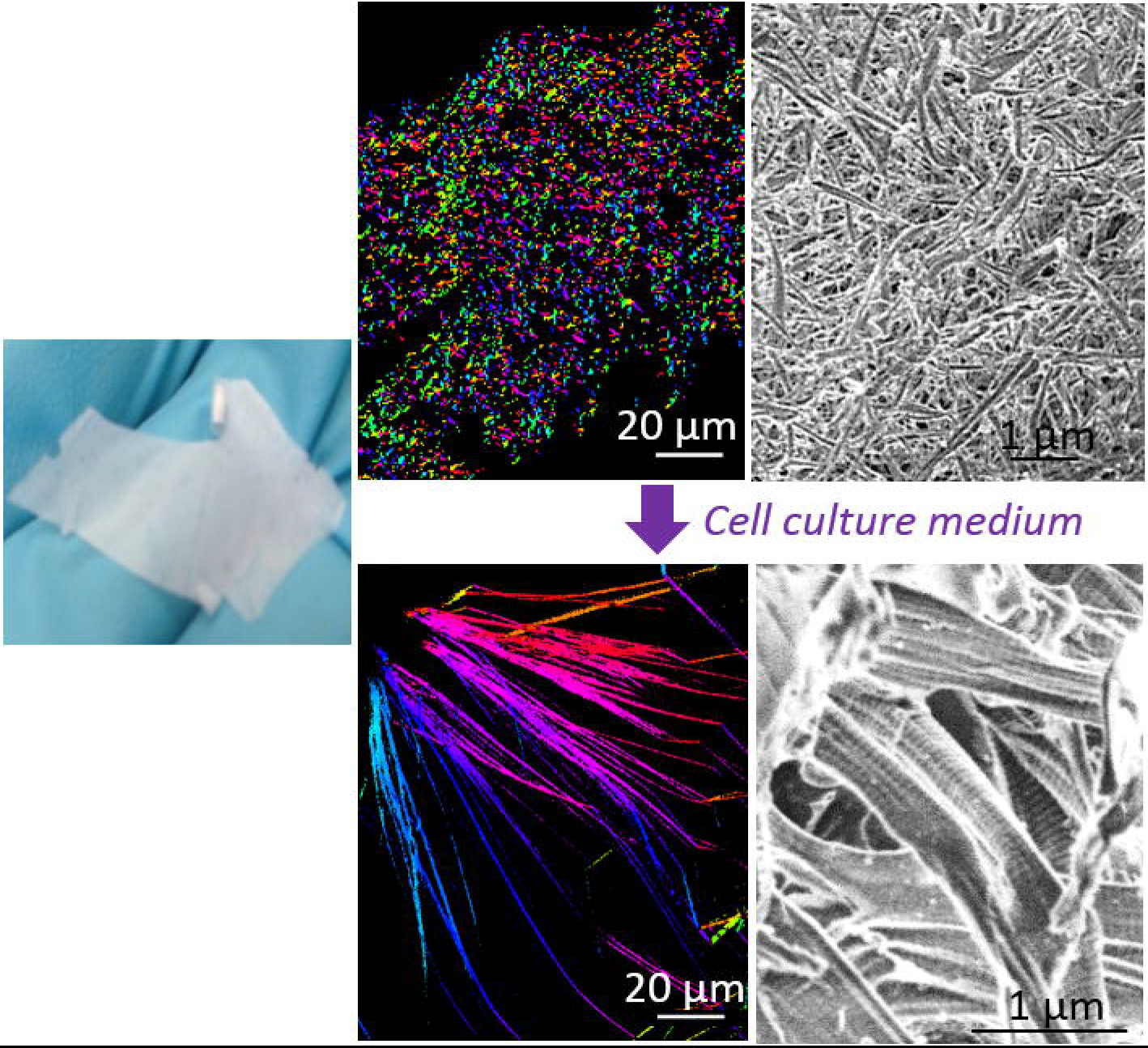

## 3. Conclusions

In conclusion, pure collagen has been successfully electrospun into a self-supported membrane, where SRCD, cryo-FE-SEM and SHG/pSHG have undoubtedly proved that collagen preserved preserves its native state triple helix structure and its ability to self-assemble into fibrils after ES. This is particularly relevant for materials coating, e.g. for prosthesis coating, with the possibility to control the thickness of the deposited membrane. In addition, we have shown the possibility to stabilize the collagen membrane by exposure to ammonia vapors, allowing to store dry membranes as pre-fibrillated scaffolds. This provides very easy to handle biomaterials of pure collagen, ready to use for biomedical applications including implantations, with preserved hierarchical structure and biological activity. Because of the final processing of ES materials into thin coatings and self-supported films, a broad range of applications can be targeted, including wound healing, cornea repair, heart patches and neural regeneration.

## 4. Methods

### 4.1. Collagen extraction and purification

Type I collagen was extracted and purified from rat tail tendons as previously described by substituting 500 mM acetic acid by 3 mM hydrochloric acid.^25,31^ Collagen purity was assessed by electrophoresis and its concentration estimated by hydroxyproline titration.^32^ All other chemicals were purchased and used as received. Water was purified with a Direct-Q system (Millipore Co.).

### 4.2. Collagen electrospinning

1.2 mL of the solution was electrospun for 4 hours at ca. 12 kV with a flow rate of 0.005 mL.min-1 in a home-made chamber flushed with dry air (ca. 10% humidity) at room temperature (~20°C). The counter electrode was a copper tape (2.5 * 1.5 cm), the distance between the needle and the counter electrode was 9 cm. After processing, the electrospun membrane was detached from the support, and a homogeneous self-standing membrane was obtained.

### 4.3. Pre-fibrillation of electrospun membranes by ammonia exposure

4 mL of ammonium hydroxide solution (25%, Carlo Erba) in a glass beaker were placed in a desiccator (153*191 mm). After 20 min exposure to ammonia vapors, collagen membranes were rinsed three times a day for 10 days in culture medium to remove all ammonia residues before cell proliferation testing. Preliminary experiments in deionized water were run to check for the decrease in pH along these washing steps.

### 4.4. Synchrotron-radiation circular dichroism

25 µL of collagen solutions in 3 mM HCl were loaded in quartz cells (Hellma) with a path length of 0.01 cm. The measurements were performed on the DISCO Beamline at SOLEIL synchrotron (Saint Aubin, France).^33^ Melting temperatures were obtained by increasing temperature from 20 to 71°C with steps of 3°C and an equilibration time of 5 min. Raw spectra were acquired from 320 to 170 nm with a 1 nm spectral resolution. Spectra were treated with the CDTool software.^34^ They are the average of three spectra. A background (HCl 3 mM acquired in the same conditions) was subtracted from them. Intensity calibration was obtained with a CSA sample.

### 4.5. Second harmonic generation

We used a custom-built laser-scanning multiphoton microscope as previously described.30 Excitation was provided by a femtosecond titanium–sapphire laser (Mai-Tai, Spectra-Physics) tuned to 860 nm, scanned in the XY directions using galvanometric mirrors and focused using a 25× objective lens (XLPLN25XWMP2, Olympus), with a resolution of typically 0.4 μm (lateral) × 1.7 μm (axial). We used either circular polarization in order to image all structures independently of their orientation in the image plane, or a set of linear polarizations with different orientations in order to perform polarization-resolved measurements. All polarization-resolved images were acquired at 36 excitation angles θ regularly spaced between 0° and 360°, using 200 kHz acquisition rate and 300×300 nm^2^ pixel size. Power at the sample was typically 12 mW for dry and stabilized membranes, and 2 mW for the hydrated samples. Epi-detection was used for dry and stabilized membranes and 2×2 binning was applied before pSHG processing. Transdetection was used for hydrated sample, with no binning. pSHG images were processed as already described^30^ to provide two images: (i) the average of all images acquired with the set of linear polarizations, that is similar to an SHG image acquired with circular polarization; (ii) a map of the main orientation of collagen in the image plane. The latter pSHG image is displayed using the HSV Look-up-table, where H is the orientation displayed in the insert, and V is the brightness, which is set to 1 if the pSHG processing is satisfactory (R2>0.5). Three areas were observed on each sample to check for the sample structure homogeneity. Each sample was investigated in triplicate.

### 4.6. SEM observations

SEM imaging was performed using a Variable Pressure Hitachi S-3400N working at an accelerating voltage of 10 kV, using an in lens secondary electron (SE) detector, with a working distance at / around 4.2 mm. Self-supported membranes were deposited on carbon-tape coated aluminum pads. Samples were coated with a 10 nm gold layer before observations.

### 4.7. Cryo-FE-SEM observations

Sample preparation and observations were performed at the Electron Microscopy Facility (EMF) of the Institut de Biologie Paris-Seine (Sorbonne University, Paris). Prior to observations, samples were sandwiched between two cupules then cryo fixed by immersion in liquid ethane using a cryofixation station (CPC, Leica). Freeze-fracture was processed at −150°C using a high vacuum station (ACE 600, Leica). FE-SEM observations were performed at −120°C at low voltage (0.790 kV) using an in lens secondary electron (SE) detector, with a 20 µm objective aperture diameter and a working distance at / around 2.5 mm (GeminiSEM 500, Zeiss; Cryo station VCT100, Leica). A sublimation step at −90°C was applied for 15 min in the SEM chamber to remove ice. Scan speed and line or drift compensation integrations were adjusted during observations.

### 4.8. Cell culture experiments

NHDF cells were cultured in complete cell culture medium: Dulbecco’s Modified Eagle’s Medium (DMEM) supplemented with 10% fetal serum, 100 U.mL-1 penicillin, 100 μg.mL-1 streptomycin, 0.25 μg.mL-1 Fungizone, and glutamax. Tissue culture flasks (75 cm2) were kept at 37°C in a 95% air : 5% CO2 atmosphere. Before confluence, cells were removed from culture flasks by treatment with 0.1% trypsin and 0.02% EDTA. Cells were rinsed and suspended in complete culture medium before use.

After pre-fibrillation of the collagen membranes, NHDF cells were cultured for 24 hours on the surface of membranes at a cell seeding density of 2 kCells / mm^2^. Cultures were incubated at 37°C in a 95% air : 5% CO2 atmosphere. Cell culture experiments were performed three times in triplicate (n=9).

### 4.9. Fluorescence microscopy (evaluation of cytocompatibility)

Collagen membranes were fixed with 4% paraformaldehyde in PBS for 1 hour, rinsed three times with PBS for 5 min each and incubated with PBS 0.2% Triton for 20 min. They were then incubated with Alexa Fluor 488 Phalloidin (Molecular Probe®) diluted 1/200 (v/v) in PBS in a dark chamber for 45 min. After rinsing three times for 5 min each in PBS, the samples were incubated with DAPI (Molecular Probe®) diluted 1/50,000 in PBS in a dark chamber for 10 min. Finally, the matrices were washed three times with PBS and mounted between slides and coverslips with the antifading solution AF3 (Biovalley) for fluorescence microscopy (ZEISS Axioplan microscope). Fibroblasts were observed at magnification ×10 and ×40.

## 5. Data availability

All data supporting the findings of this study are available from the corresponding authors upon request.

## Supporting information

Supplementary Information: SEM of collagen ES during optimization; Macroscopic structure of the fibrillary self-supported membrane

## 6. Acknowledgements

We thank Gervaise Mosser, Alexandre Bahezre, for helpful discussions; Antoine Fayssinet for experimental help; Ghislaine Frébourg, Virginie Bazin, Alexis Canette, Michaël Trichet for cryo-FE-SEM observations performed at the Electron Microscopy Facility (EMF) of the Institut de Biologie Paris-Seine (FR3631, Sorbonne University, CNRS, Paris). SRCD on DISCO beamline at the SOLEIL Synchrotron was performed under proposals 20180440 and 20170028. CA thanks Florence Babonneau for her constant support.

