## Supplementary Information: SEM of collagen ES during optimization; Macroscopic structure of the fibrillary self-supported membrane for "Electrospinning of pure, native, cross-linker free self-supported collagen membrane"

### 1. Optimization of the solution conditions to improve collagen electrospinnability

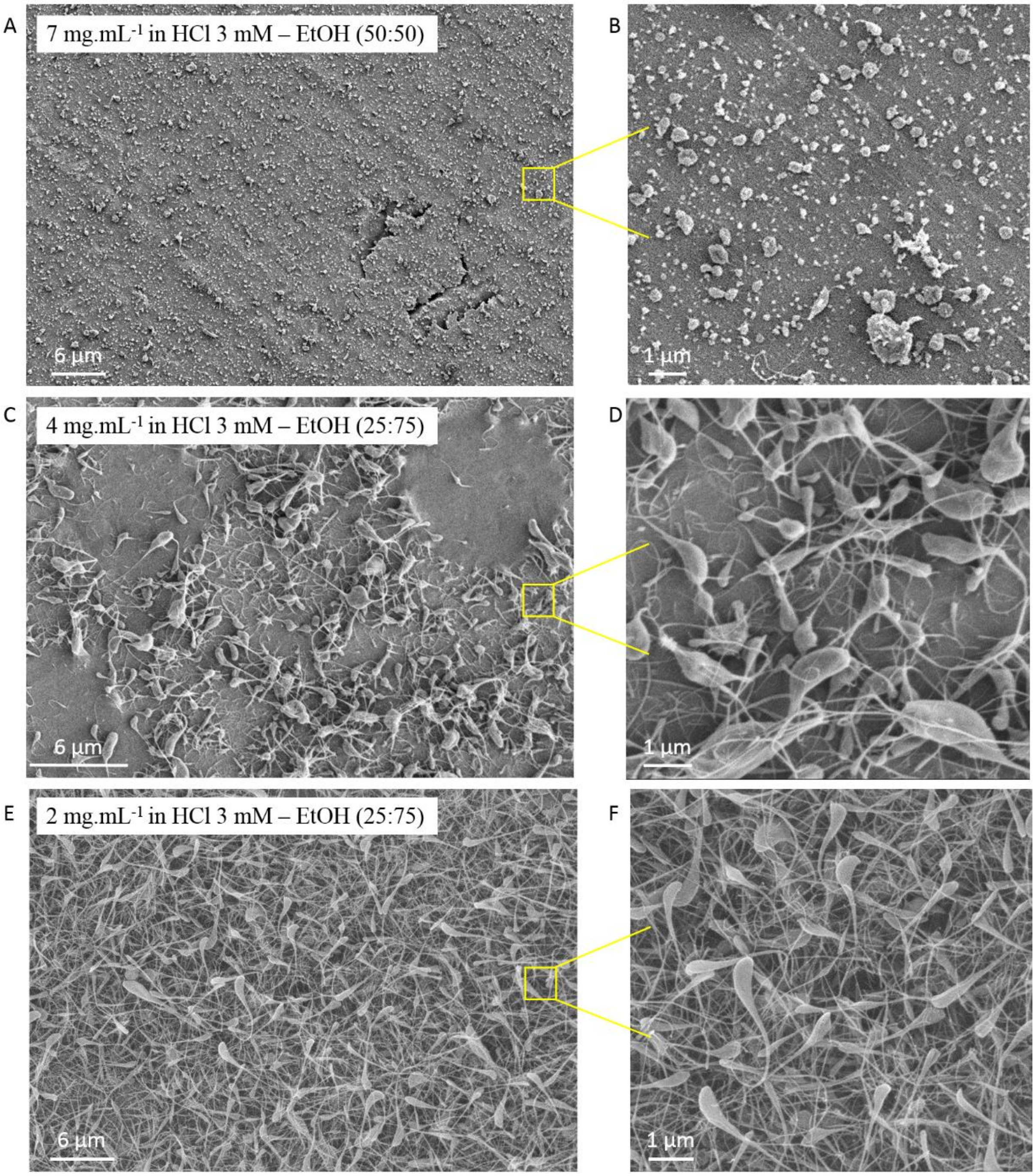


**Figure S1.** SEM photos of collagen electrospinning during optimization of the solution conditions (solvent and collagen concentration).

### 2. Macroscopic structure of the fibrillary self-supported membrane

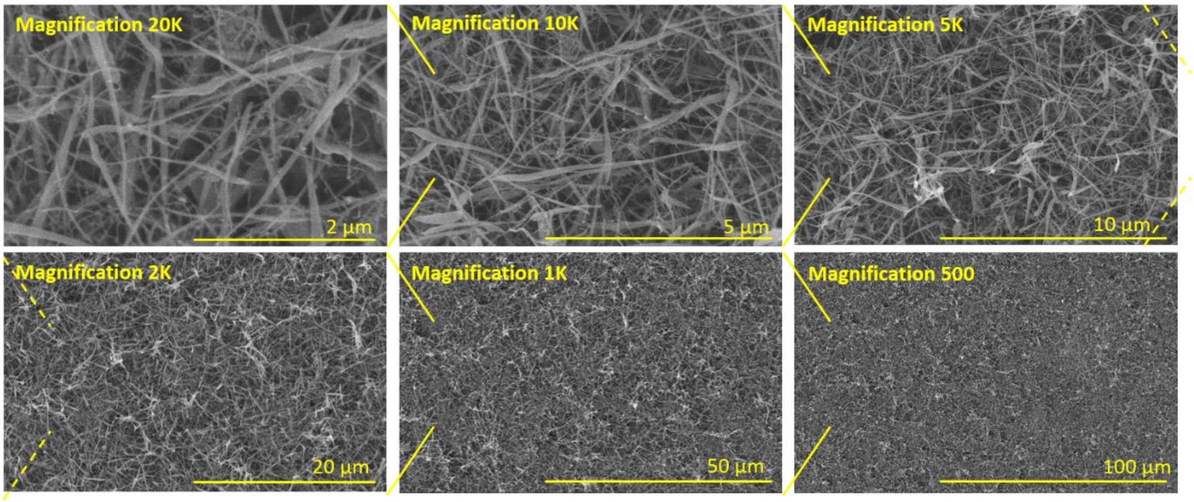


**Fig. S2.** SEM images of a self-supported collagen membrane obtained by ES ([collagen] = 1.7 $mg.mL^{-1}$ in 30 mM HCl:EtOH (25:75 v/v)) at different magnifications (from 20K to 500) to highlight the macroscopic structure of the fibrillary self-supported membrane.
